## Supplementary Material for "Microbiota-directed fibre activates both targeted and secondary metabolic shifts in the distal gut"

This PDF file includes:

I. Supplementary Figures and Legends
II. Supplementary Tables and Legends
III. Supplementary Datasets and Legends
IV. References cited in the Supplementary Material

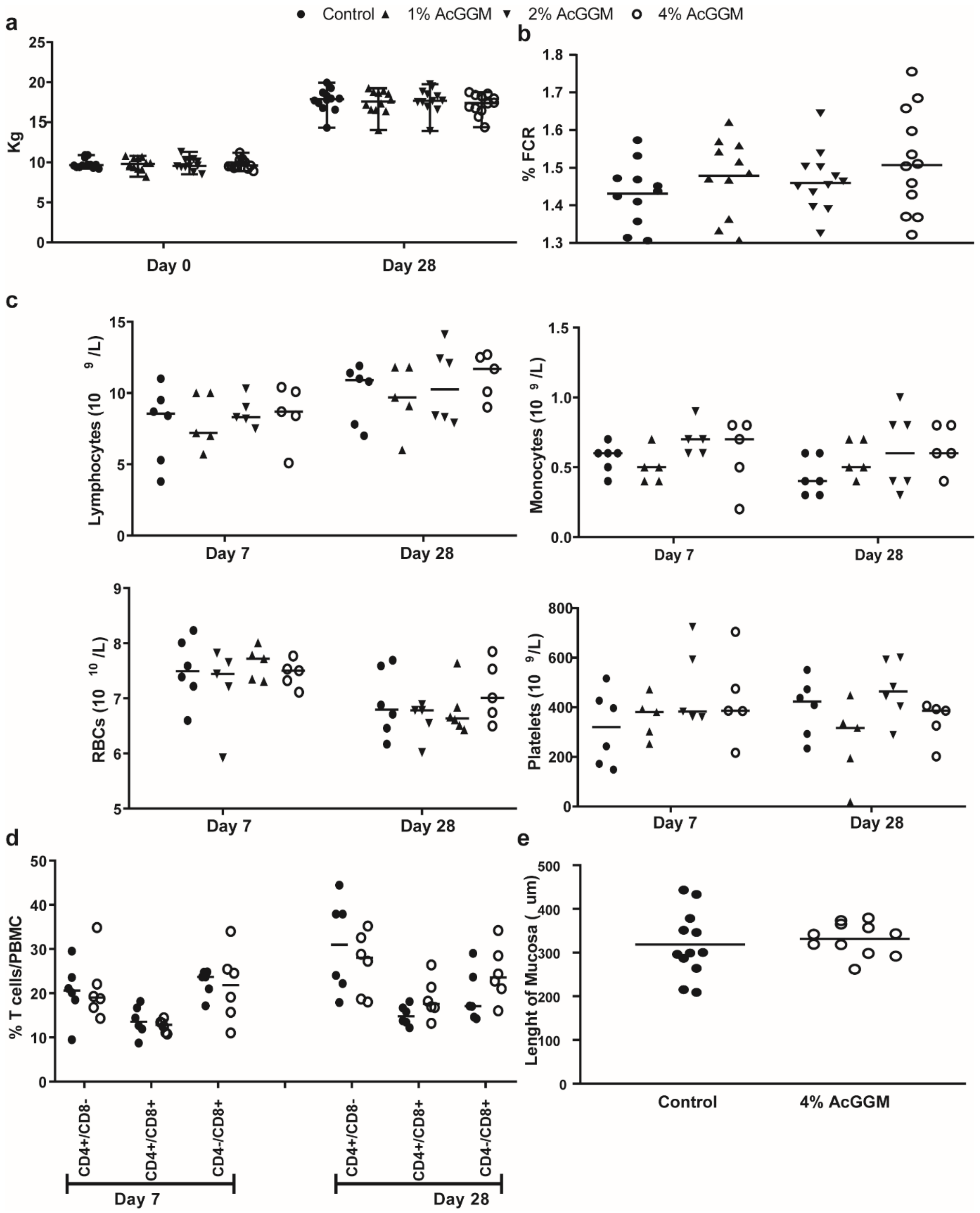


**Supplementary Figure 1. Performance metrics and health status of weaning piglets subjected to various AcGGM-containing feeds. a.** Average weight of piglets at the start of the feeding trial and at sampling day shows no statistically significant difference in Dunnett's multiple comparisons test (p=0.05). **b.** Feed conversion rate for piglets in the 4% AcGGM inclusion levels shows no statistically significant difference Dunnett's multiple comparisons test (p=0.05). **c.** Neither the numbers of lymphocytes, RBCs, monocytes nor platelets were significantly affected by the inclusion of AcGGM in diet (Tukey's multiple comparisons, p=0.05). **d.** Flow cytometry analysis of T cell populations in piglets from the control vs 4%

AcGGM inclusion group shows no statistically significant shift in any of the subpopulations (two-way ANOVA, p=0.05) (gating strategy: see Supplementary Figure 6). Y-axis - abundance of cells in percent of peripheral blood mononuclear cells. **e.** Colon epithelium morphology: length of mucosal layer shows no significant difference between the controls and 4% AcGGM inclusion. Numerical values shown in the charts are available in Supplementary Table 1.

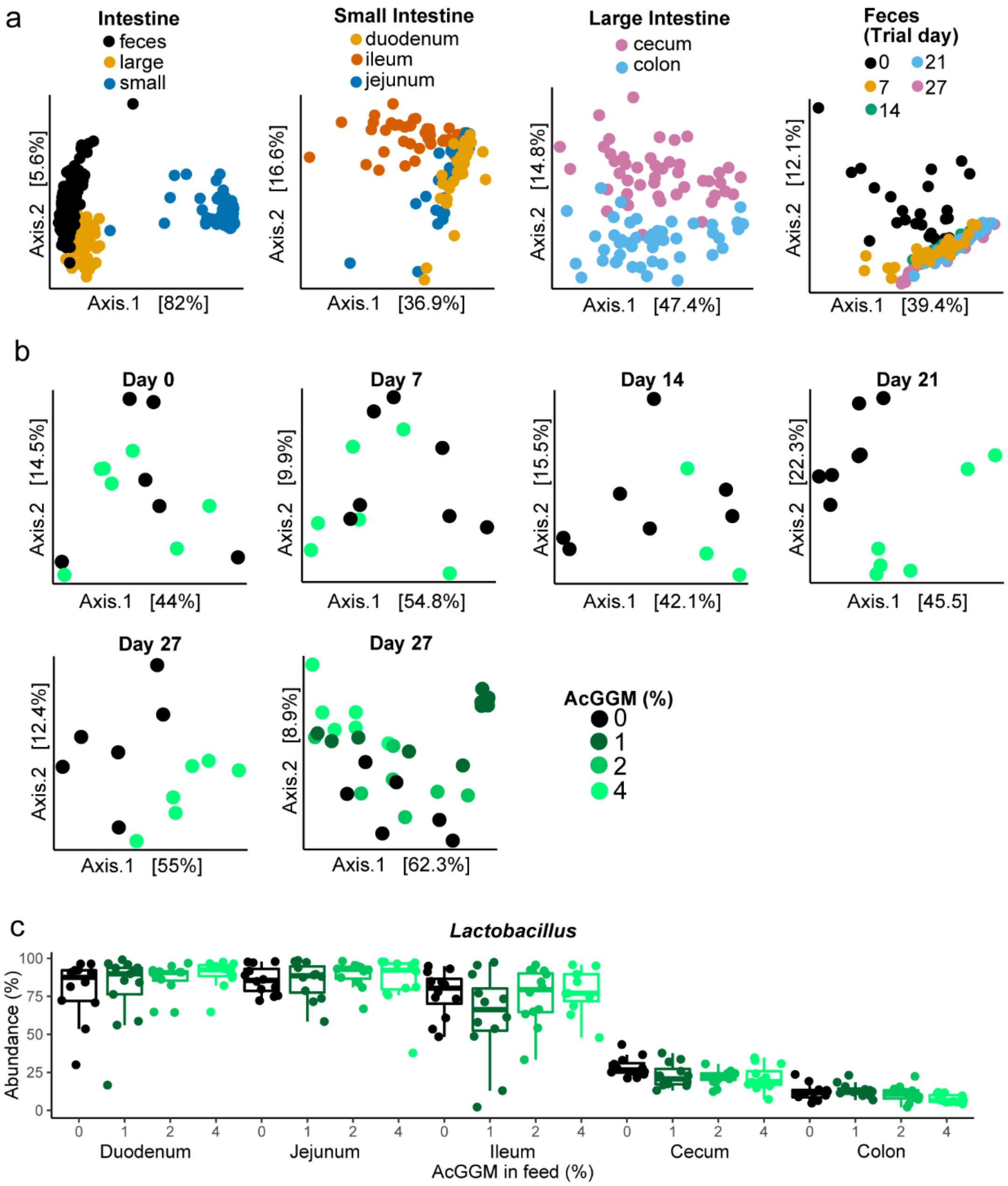


**Supplementary Figure 2. Spatial and temporal changes in the gut microbiome of weaning piglets fed varying levels of AcGGM**. Structural microbiome changes were monitored using DNA extracted from gut or fecal samples and 16S rRNA gene analysis over the month-long trial. **a.** Spatial variation of the gut microbiome of weaning piglets was observed between different regions within the small and large intestines as well as feces. **b.** Temporal microbiome changes of weaning piglets fed varying levels of AcGGM. Ordination plots of Bray-Curtis distances between microbiome communities analyzed from fecal samples collected from 6 randomly selected animals per treatment group throughout the 28 day trial. From day 14 onwards, greater variation was observed between the control samples and the samples from the three AcGGM inclusion levels (1, 2 and 4%), indicating structural changes in the pig fecal microbiome composition in response to AcGGM inclusion. **c.** Spatial variation was observed for specific taxa such as the *Lactobacillus*, which was observed at higher relative abundances in regions within the small intestine. Bars represent median and boxes interquartile range determined from 12 animals analyzed per dietary group.

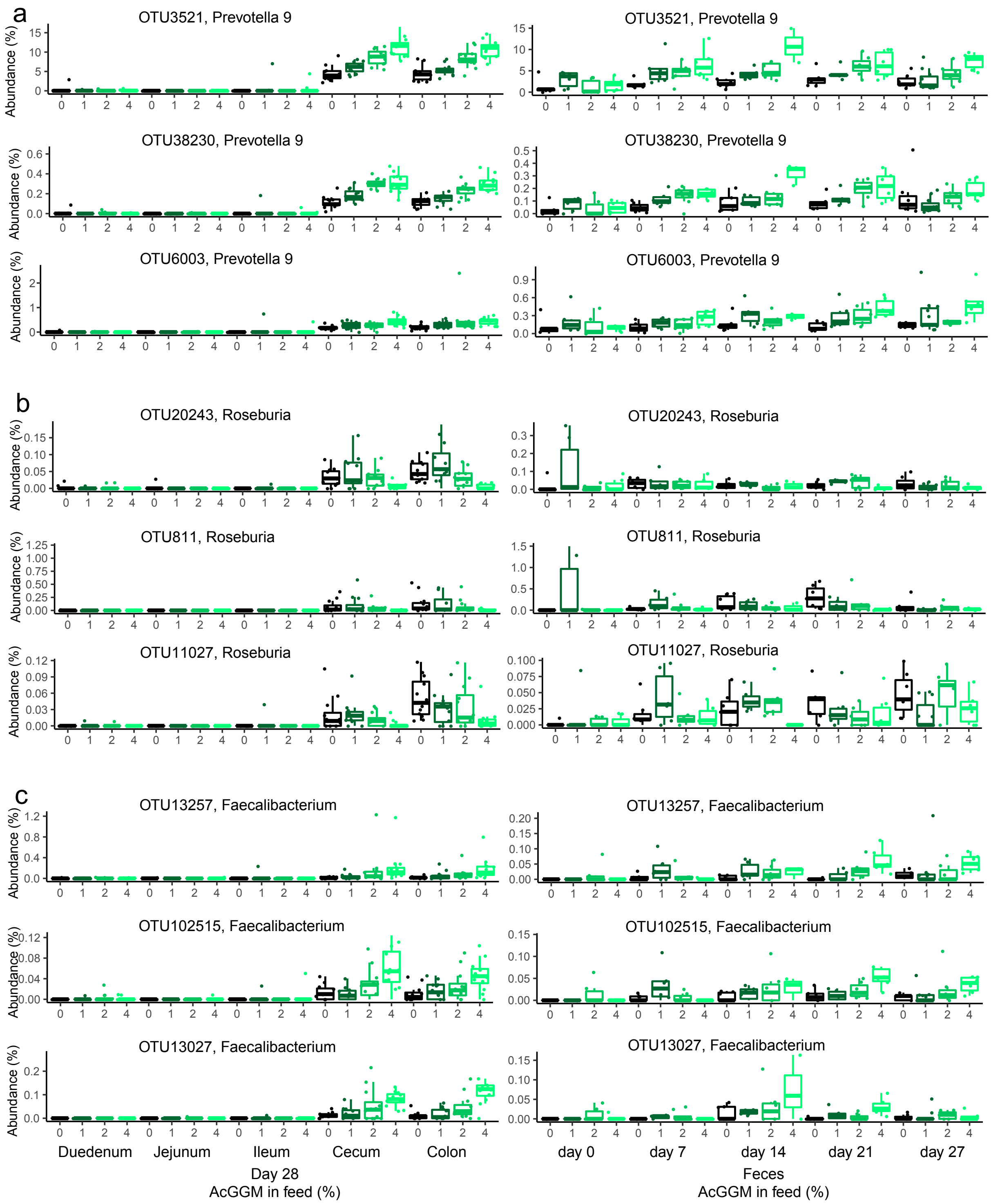


**Supplementary Figure 3. Spatial and temporal changes of selected *Prevotella*- (a), *Roseburia*- (b) and *Faecalibacterium*- (c) affiliated phylotypes in fecal microbiome from weaning piglets fed varying levels of AcGGM.** The list of OTU abundances that changed significantly (adjusted $p<0.05$) in response to AcGGM diets are listed in Supplementary Dataset 2. Bars represent median and boxes interquartile range determined from 12 animals analyzed per dietary group.

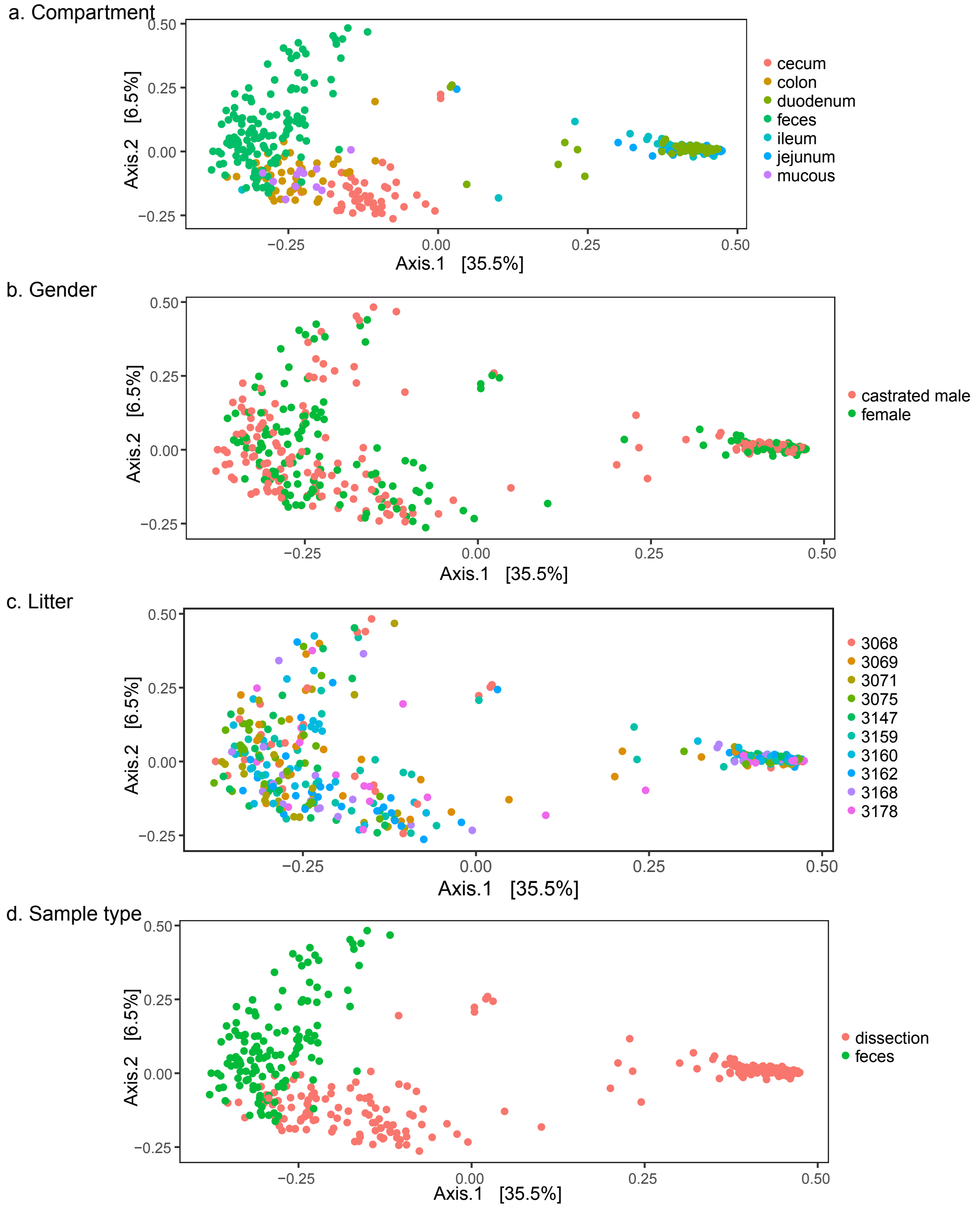


**Supplementary Figure 4. Microbiota composition of samples from different sites in the pig gastrointestinal tract (GIT).** Data is presented as a multidimensional scaling (MDS) ordination of weighted UNIFRAC distances. **a** Samples from all GIT compartments (including feces), were labelled according to GIT compartment of origin. The two clusters separate samples from upper GIT (duodenum, jejunum, ileum) in the right side of the plot, from fecal samples and lower GIT and (colon, cecum, colon mucus). No biases were observed from any of the variables (**b** Piglet gender. **c** Litter **d** Sample type) listed here.

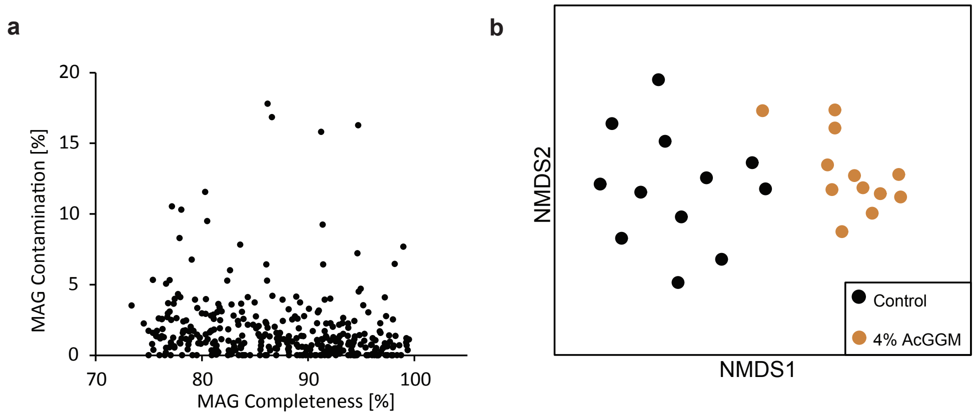


**Supplementary Figure 5. Metagenomic and metaproteomic analysis of the colon microbiome population of control and 4% AcGGM fed piglets. a.** Completeness % vs contamination % of the 355 MAGs recovered from 24 colon samples. Completeness and contamination was determined for each MAG using CheckM[1] version 1.0.7. A total of 145 had >90% completeness and were considered high quality according to the Genomics Consortium Standards[2] **b.** Non-metric multidimensional scaling (NMDS) ordination plot of sample distances calculated with MASH based analysis on the genetic content of MAGs recovered from metagenomes, sequenced and analyzed from colon samples collected from animals fed the control (n=12) and 4% AcGGM diets (n=12). Sample distribution shows the piglets from each dietary group had distinct genomic composition.

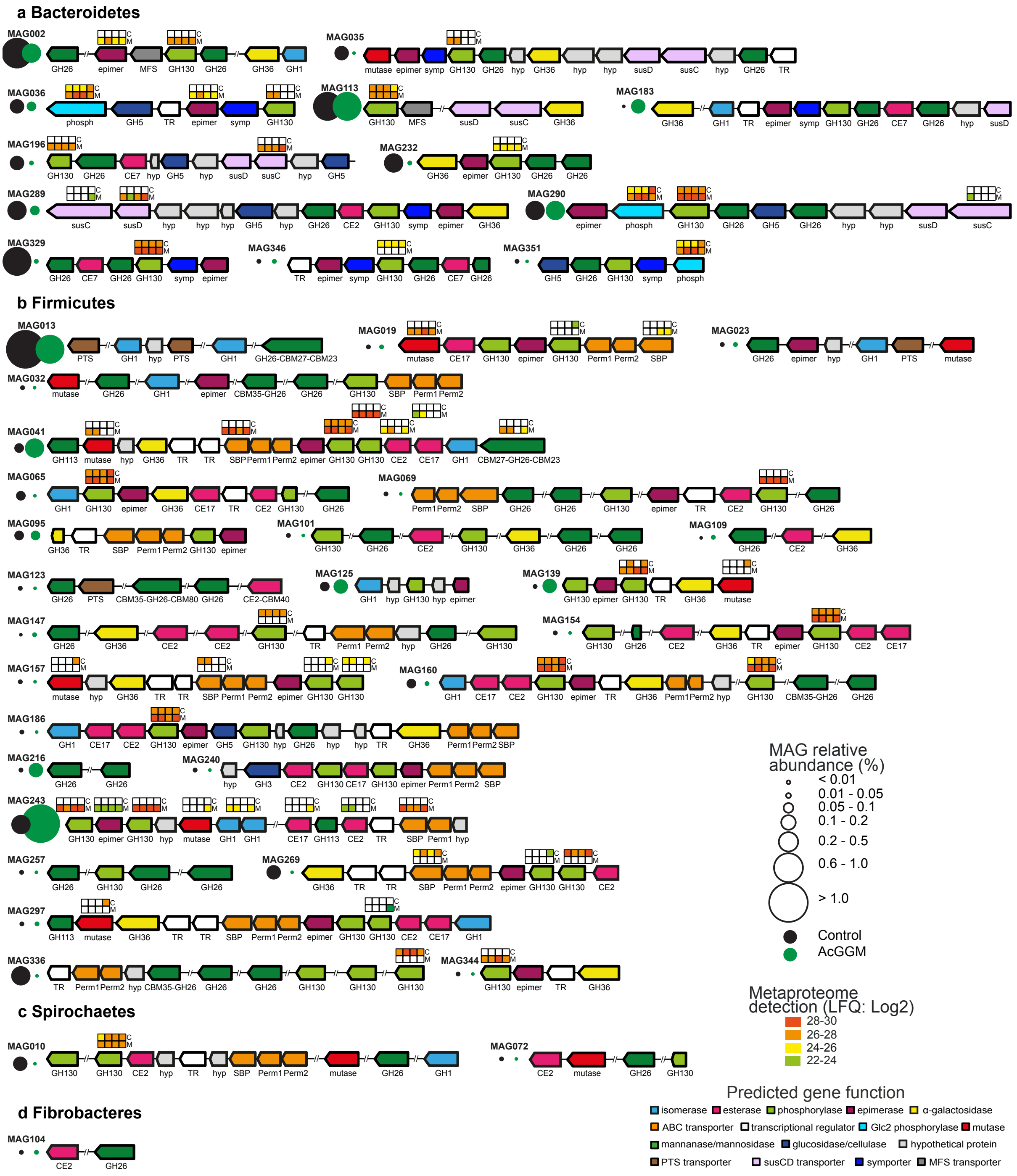


**Supplementary Figure 6. Metaproteomic detection of predicted mannan-PULs encoded in Firmicutes-affiliated MAGs in pigs fed with the control or 4% AcGGM diet.** Predicted PULs were determined using PULDB[3] (Bacteroidetes), which combines CAZymes[4], outer-membrane transport and carbohydrate-binding lipoproteins (SusC/D-like), as well as from previous biochemical and structural characterization of the mannan degradation cluster in *R. intestinalis* L1-82[5]. Clusters were affiliated to phyla: Bacteroidetes (**a**), Firmicutes (**b**), Spirochaetes (**c**) and Fibrobacteres (**d**). Heat maps above detected enzymes show the LFQ detection levels for the four replicates sampled in control (C) and 4% AcGGM-fed (M) pigs.

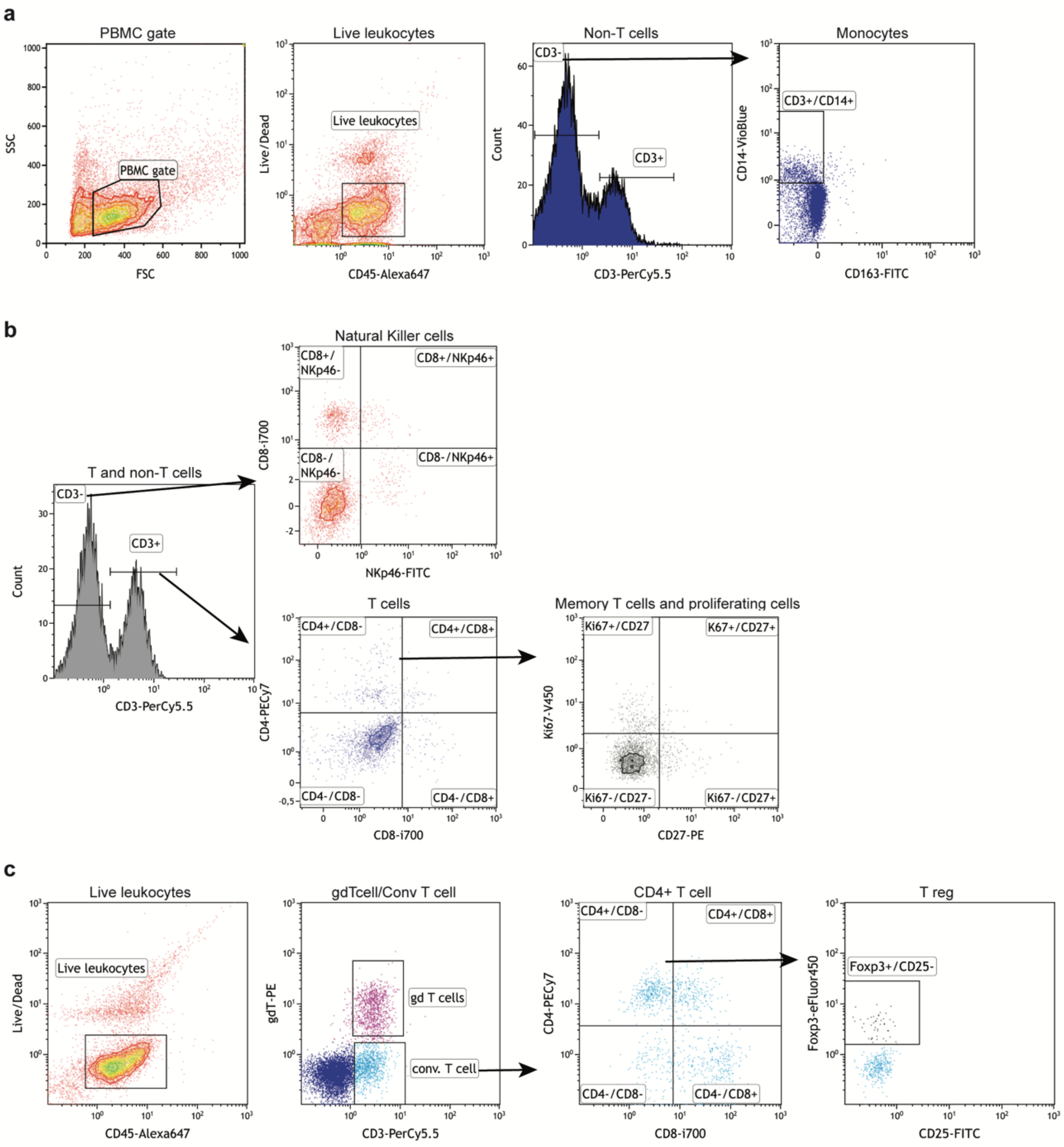


**Supplementary Figure 7. Gating strategy used for flow cytometric data presented in Supplementary Figure 1. a.** Cells were gated on SSC vs FSC to determine size and granularity, thereafter cells were gated based on live/dead to identify live leukocytes, which were used for further analysis. Monocytes were assigned as CD45+, CD3- and CD14+ or CD163+. **b.** Natural killer cells were assigned as CD45+, CD3- and CD8+ or NKp46+ while the different subsets of T cells were determined according to the expression of CD4, CD8, Ki67 and CD27. **c.** T reg were identified according to the expression of gdT (gdT cells) or CD3, CD4, CD8 (conventional T cells) or Foxp3 and CD25 (T reg). Supplementary Table 2 outlines the percentage of each gate.

**Supplementary Table 1**. **Evaluation of colon morphology**. Analysis was performed according to the scoring system described by Fabia et al.[6] and indicates no significant difference between the control and 4% AcGGM inclusion level.

| | 0% AcGGM | 4% AcGGM |
|---|---|---|
| **Ulceration** | 0 | 0 |
| **Mucosal atrophy** | 0 | 0 |
| **Edema** | 0 | 0 |
| **Cell infiltration** | 0.6 | 0.5 |
| **Vascular dilation** | 0.8 | 0.3 |
| **Length of mucosa (µm)** | 318±21.6 | 332±11.2 |

**Supplementary Table 2**. Total reads and sequence length obtained in the whole metagenome sequencing of each sample. 0M and 4M in "Sample id" designate 0% and 4% AcGGM content in diet.

| Sample id | total reads per sample | total bases per sample | total Gbases per sample | Accession number (SRA) |
|---|---|---|---|---|
| **01_0M** | 149875718 | 22481357700 | 22.48 | SRR10209687 |
| **02_0M** | 39328376 | 5899256400 | 5.89 | SRR10209686 |
| **03_0M** | 41490606 | 6223590900 | 6.22 | SRR10209675 |
| **04_0M** | 50328468 | 7549270200 | 7.54 | SRR10209670 |
| **05_0M** | 51520620 | 7728093000 | 7.72 | SRR10209669 |
| **06_0M** | 44035782 | 6605367300 | 6.60 | SRR10209668 |
| **07_0M** | 71111130 | 10666669500 | 10.66 | SRR10209667 |
| **08_0M** | 38528542 | 5779281300 | 5.78 | SRR10209666 |
| **09_0M** | 48341876 | 7251281400 | 7.25 | SRR10209665 |
| **10_0M** | 43270632 | 6490594800 | 6.50 | SRR10209664 |
| **11_0M** | 86191960 | 12928794000 | 12.93 | SRR10209685 |
| **12_0M** | 60690558 | 9103583700 | 9.10 | SRR10209684 |
| | | **0M total:** | **108.71** | |
| **13_4M** | 176308520 | 26446278000 | 26.45 | SRR10209683 |
| **14_4M** | 46804332 | 7020649800 | 7.02 | SRR10209682 |
| **15_4M** | 53833158 | 8074973700 | 8.07 | SRR10209681 |
| **16_4M** | 49041518 | 7356227700 | 7.35 | SRR10209680 |
| **17_4M** | 61654754 | 9248213100 | 9.24 | SRR10209679 |
| **18_4M** | 42908204 | 6436230600 | 6.43 | SRR10209678 |
| **19_4M** | 34987350 | 5248102500 | 5.25 | SRR10209677 |
| **20_4M** | 44732402 | 6709860300 | 6.71 | SRR10209676 |
| **21_4M** | 33761876 | 5064281400 | 5.06 | SRR10209674 |
| **22_4M** | 55012540 | 8251881000 | 8.25 | SRR10209673 |
| **23_4M** | 45067490 | 6760123500 | 6.76 | SRR10209672 |
| **24_4M** | 40208238 | 6031235700 | 6.03 | SRR10209671 |
| | | **4M total:** | **102.65** | |
| | | mean: | 8.8 | |
| | | median: | 7.1 | |
| | | max: | 26.4 | |
| | | min: | 5.1 | |

**Supplementary Table 3. Composition and chemical content of basal diet.**

| Ingredients (%) | | Calculated content (g/Kg) | |
|---|---|---|---|
| Wheat | 51 | Energy, MJ/kg | 14.14 |
| Barley | 20 | Dry Matter | 884 |
| Soybean meal | 8 | Crude protein | 176.91 |
| Oats | 6 | Crude fibre | 34 |
| Soy oil | 4 | Digestible crude protein | 156.11 |
| Fish meal | 2 | Starch | 431.28 |
| Potato protein | 2 | Crude fat | 55 |
| Corn gluten | 1.78 | Calcium | 9.13 |
| Calcium phosphate | 1.13 | Phosphorous | 5.82 |
| Limestone | 1 | Digestible phosphorous | 3.73 |
| Vitamin mix | 1 | Sodium | 2.51 |
| Lysine | 0.69 | Chloride | 4.59 |
| Salt | 0.39 | Lysine | 13.04 |
| Sodium bicarbonate | 0.36 | Digestible Lys | 12.06 |
| Threonine | 0.25 | Methionine + Cysteine | 7.27 |
| Methionine | 0.15 | Digestible Methionine + Cysteine | 6.57 |
| Valine | 0.14 | Methionine | 4.21 |
| Tryptophan | 0.075 | Threonine | 8.11 |
| | | Tryptophan | 2.70 |
| | | Valine | 9.03 |

AcGGM was assigned net energy value zero.

*VilomixMineral premix and vitamin mineral premix provided the following per kilogram of diet: vitamin A, 12 000 IU; vitamin $D_3$, 3200 IU; vitamin E, 80 IU; vitamin $K_3$, 25 mg; vitamin $B_1$, 25 mg; vitamin $B_2$, 65 mg; vitamin $B_6$, 5 mg; vitamin $B_{12}$, 0.5 mg; niacin, 45 mg; pantothenic acid, 20 mg; folic acid 15 mg; biotin 0.15 mg. §Fe,150 mg; Cu, 125 mg; Zn, 150 mg; Mn, 30 mg; I, 0.3 mg; Se, 0.3 mg.

**Supplementary Dataset 1**. Metadata recorded from weaning piglets subjected to various AcGGM-containing feeds. Measurements include: weight, feed conversion ratio, digesta pH, flow cytometry, hematology and fecal score. Weight, Feed conversion ratio and digesta pH were measured for the 48 piglets, while flow cytometry was performed on 12 animals, 6 piglets control and 6 piglets 4% AcGGM. Hematology measurements were determinate for 24 animals, 6 piglets per diet. Fecal scores were determined daily per pen and were quantified according to firmness and shape as described in Material and Methods, with score 1 = firm and shaped and score = 4 watery. Samples with score 3 or 4 are considered diarrheic. SCFA content of digesta samples from cecum and colon was analyzed on an RSLC Ultimate 3000 (Dionex, USA) HPLC using a REZEX ROA-Organic Acid H+ 300x7.8mm ion exclusion column (Phenomenex, USA) at 65°C, 10 µL injection volume, with isocratic elution using 0.6 mL/min of 5mM $H_2SO_4$ as mobile phase and a UV detector set to 210 nm. The SCFAs were recorded as millimolar concentration in the digesta and analyzed using a two-tailed *t* test.

**Supplementary Dataset 2. Relative abundance of operational taxonomic units that were observed to change (adjusted p<0.05) in response to one month of the host animals being fed varying AcGGM inclusion levels.** Samples from the cecum and colon were collected at day 28 and analyzed using 16S rRNA gene sequencing analysis with the MetagenomeSeq fitZIG and DESeq2[7] negative binomial algorithms via the QIIME wrapper.

**Supplementary Dataset 3. Relative abundance and taxonomic affiliation of MAGs reconstructed from the colon of pigs fed either the control or 4% AcGGM diet.** Taxonomic classification (determined via GTDB-Tk) is given for 355 MAGs, while their relative abundance (determined via CoverM) across metagenomes generated for 12 x control pigs (01_0M-12_0M) and 12 x 4% AcGGM pigs (13_4M-24_4M) are listed and analyzed using a two-tailed *t* test.

**Supplementary Dataset 4.** Concatenated ribosomal protein tree of 22 ribosomal proteins for all MAGs in the pig distal gut microbiome, with reference sequences in Newick format.

**Supplementary Dataset 5. Total counts, means and standard deviation of detected proteins mapped to MAGs analyzed in weaning piglets fed either the control or 4% AcGGM diets.** Four randomly selected colon samples from animals fed either the control (C) or mannan (M) diet were selected for metaprotemic analysis, which detected a total of 8515 protein groups that mapped against 355 MAGs reconstructed from the colon.

**Supplementary Dataset 6. Metabolic reconstruction for selected MAGs and LFQ intensities of their detected proteins in samples analyzed from weaning piglets fed either the control (C) or 4% AcGGM (M) diet.** Key metabolic enzymes and pathways are annotated for each ORF within each MAG (E.C. and CAZy family where available) as well as gene names are provided. Number and gene names in square parenthesis indicate enzymes contributing to metabolic pathways that are illustrated in **Fig. 7**. Pathway reconstruction is provided for predicted butyrate-producers (MAG041, MAG243, MAG133, MAG269, MAG292, MAG324), *Prevotella*-affiliated populations (MAG191, MAG285, MAG196, MAG034) and other populations whose proteomes were highly detected/enriched in our analysis (MAG053, MAG150, MAG013, MAG225, MAG048). *The complete E.C. list for each CAZy family can be found at CAZy.org. In many instances, multiple E.C. numbers are listed for each CAZy family and are constantly being upgraded as more biochemical information becomes available.

**Supplementary Dataset 7. MAG enrichment analysis**. MAGs enriched in hierarchal metaproteome expression clusters (see **Fig. 4c**), determined using 4562 unique groups (consisting of 12 535 shared proteins). That is, MAGs that contribute with more detected proteins in a cluster than what we would expect by chance. Five different clusters were observed, with protein groups differentially detected in AcGGM fed pigs (M1-4), control pigs (C1-4), all pigs (M1-4 + C1-4), AcGGM fed pigs plus one control (M1-4 + C4) and only in individual pigs (Individual). MAGs in each cluster are ranked by adjusted p-value. **x** denotes total number of detected (shared) proteins for a given MAG within a given expression cluster. **k** denotes total number of detected (shared) proteins within a given expression cluster. **m** denotes total number of detected (shared) proteins within a given MAG. **N** denotes total number of detected (shared) proteins in the complete dataset

**Supplementary Dataset 8. MAPP analysis**. Micro Array Polymer Profiling (MAPP) of plant cell wall components in each diet prior to feeding as well as colon samples. Colour intensity is proportional to mean spot signal. Starch was the most detected fibre in the basal feed and was readily digested in pigs fed either the control or AcGGM diets. Mannan fibres in both the basal feed as well as AcGGM were not detected, presumably due to the lack of specificity between the *Thermotoga maritima* CBM27 probe (Genbank: AAD36302) and/or the ineffectiveness of the extraction method to dissociate mannan fibres from other plant cell wall fibres in order for them to be detected. Xylan and xyloglucan fibres were observed to increase in digesta samples, presumably due to the deconstruction of plant cell walls and the liberation of these fibres.

**Supplementary Dataset 9. EC enrichment analysis**. EC-annotations enriched in hierarchal metaproteome expression clusters (see **Fig. 4c**), determined using 4562 unique groups (consisting of 12 535 shared proteins). That is, designated EC annotations that contribute with more detected proteins in a cluster than what we would expect by chance. Five different clusters were observed, with protein groups differentially detected in AcGGM fed pigs (M1-4), control pigs (C1-4), all pigs (M1-4 + C1-4), AcGGM fed pigs plus one control (M1-4 + C4) and only in individual pigs (Individual). EC-annotations in each cluster are ranked by adjusted p-value. **x** denotes total number of detected protein groups for a given EC annotation number within a given expression cluster. **k** denotes total number of detected (shared) proteins within a given expression cluster with an EC annotation number. **m** denotes total number of detected protein groups with a given EC annotation number. **N** denotes total number of detected protein groups in the complete dataset with an EC annotation number.

**REFERENCES.**


1 Parks, D. H., Imelfort, M., Skennerton, C. T., Hugenholtz, P. & Tyson, G. W. CheckM: assessing the quality of microbial genomes recovered from isolates, single cells, and metagenomes. *Genome Res* **25** (2015).

2 Bowers, R. M. *et al.* Minimum information about a single amplified genome (MISAG) and a metagenome-assembled genome (MIMAG) of bacteria and archaea. *Nat Biotechnol.* **35**, 725-731 (2017).

3 Terrapon, N., Lombard, V., Gilbert, H. J. & Henrissat, B. Automatic Prediction of Polysaccharide Utilization Loci in Bacteroidetes Species. *Bioinformatics* **31**, 647-655 (2014).

4 Martens, E. C., Koropatkin, N. M., Smith, T. J. & Gordon, J. I. Complex glycan catabolism by the human gut microbiota: the Bacteroidetes Sus-like paradigm. *J. Biol. Chem.* **284**, 24673-24677 (2009).

5 La Rosa, S. L. *et al.* The human gut Firmicute *Roseburia intestinalis* is a primary degrader of dietary β-mannans. *Nat. Commun.* **10**, 905 (2019).

6 Fabia, R. *et al.* The Effect of Exogenous Administration of Lactobacillus reuteri R2LC and Oat Fiber on Acetic Acid-Induced Colitis in the Rat. *Scandinavian Journal of Gastroenterology* **28**, 155-162 (1993).

7 Love, M. I., Huber, W. & Anders, S. Moderated estimation of fold change and dispersion for RNA-seq data with DESeq2. *Genome Biol.* **15**, 550 (2014).
